# Correcting overprediction reduces the propagation of uncertainty from species distribution models into spatial conservation prioritization

**DOI:** 10.64898/2026.05.19.726420

**Authors:** Thiago Cavalcante, Sara Si-Moussi, Marianne Tzivanopoulos, Maxime Hoareau, Wilfried Thuiller, Heini Kujala

## Abstract

Effective conservation planning increasingly relies on species distribution models (SDMs) to guide where actions deliver the greatest biodiversity benefits through spatial conservation prioritization. However, SDMs are inherently uncertain, and this uncertainty propagates through prioritization processes, affecting the identification of priority areas and influencing conservation decisions. Here, we evaluate whether correcting SDM overprediction reduces uncertainty propagation into spatial conservation prioritization. Using two large European datasets of vertebrates and invertebrates, we compared unconstrained SDMs with models corrected for overprediction through a Bayesian integration of occurrences, expert range maps, and habitat suitability. We found that overprediction correction reduced spatial and performance uncertainty, with uncertainty strongly structured by model and algorithm choice and amplified when overprediction was not corrected. Although no single modelling adjustment fully eliminates uncertainty propagation from SDMs into prioritization, we demonstrate that overprediction correction consistently reduces it across datasets, taxa, and modelling approaches, highlighting its importance for robust conservation planning.

## Introduction

Effective conservation depends on identifying areas where actions will deliver the greatest benefits to biodiversity, a task often complicated by incomplete knowledge of species distributions and ecosystem dynamics (Whittaker et al., 2005; Cardoso et al., 2011; Moilanen, 2012; Diniz-Filho et al., 2013; Giakoumi et al., 2025). Species distribution models (SDMs) play a central role in this process by predicting where species are likely to occur based on environmental conditions and observed occurrences (Thuiller, 2024), thereby providing key inputs for conservation planning across regions and taxa (Guisan et al., 2013). To translate these predictions into actionable guidance, SDM outputs are incorporated into spatial conservation prioritization (SCP), a methodological framework that enables the systematic identification of areas that maximize benefits for biodiversity goals (Margules and Pressey, 2000; Wilson et al., 2009; Giakoumi et al., 2025). These approaches increasingly rely on SDM outputs to rank areas by their contribution to biodiversity and to guide decisions such as protected area expansion, restoration, and land-use planning.

As these model-based approaches play an expanding role in real-world conservation practice, their reliability becomes critically important. However, SDMs are inherently uncertain (Thuiller et al., 2019), and this uncertainty can propagate through prioritization processes, affecting both the identification of priority areas and the expected representation of species (Beale and Lennon, 2012; Guisan et al., 2013; Meller et al., 2014; Velazco et al., 2020; Muscatello et al., 2021). Understanding this uncertainty and implementing measures to buffer its effects is therefore essential for robust and reliable conservation decision-making.

Among the different sources of uncertainty in SDMs, overprediction—where models assign suitable conditions to areas outside the true distribution of a species—is particularly pervasive (Soberon and Peterson, 2005; Mendes et al., 2020). Overprediction can arise from model extrapolation, incomplete sampling, limitations in environmental predictors or the neglect of dispersal processes, and tends to inflate estimates of species ranges. In the context of spatial conservation prioritization, this bias can lead to the allocation of conservation value to areas where species are unlikely to occur, potentially misdirecting conservation efforts (Velazco et al., 2020).

Efforts to address overprediction have shown that constraining SDMs (e.g., by incorporating biogeographical information) improves the spatial accuracy of the models and consequently the reliability of conservation prioritization (Loiselle et al., 2003; Hannemann et al., 2016; Mendes et al., 2020; Velazco et al., 2020). However, previous studies have generally focused on averaged predictions or relied on a single modelling algorithm, without examining how overprediction interacts with the uncertainty arising from different modelling algorithms and ensemble approaches. As a result, it remains unclear whether correcting overprediction consistently improves prioritization outcomes when accounting for variability across modelling choices. Understanding these effects is therefore essential for producing more reliable and resilient conservation plans.

In this study, we assess whether correcting overprediction reduces the propagation of uncertainty from SDMs into spatial conservation prioritization. We explicitly quantify how overprediction influences uncertainty across multiple modelling algorithms and ensemble approaches, using both spatial priority rankings and performance curves as complementary measures of prioritization outcomes. By comparing constrained and unconstrained SDM scenarios across taxa and modelling strategies, we provide a systematic evaluation of the role of overprediction in shaping both the magnitude and spatial structure of uncertainty. Our results offer new insights into how improving SDM realism can enhance the robustness and reliability of conservation planning.

## Methods

### Species distribution models and overprediction correction

We analysed two independent datasets of terrestrial vertebrates and invertebrates across Europe at 1km^2^ resolution (see Si-Moussi et al., 2025 for methodological details). The vertebrate dataset comprised 1,050 species across amphibians, birds, mammals, and reptiles, with SDMs based on Random Forest (RF), extreme gradient boosting (XGBoost), and multi-layer perceptron neural networks (MLP). The invertebrate dataset included 64 Habitats Directive species across seven taxonomic groups (Araneae, Carabidae, Gastropoda, Heterocera, Odonata, Orthoptera, and Rhopalocera), with SDMs generated using GLM, GAM, Random Forest (RF), and XGBoost (see Supporting Information S1 for further details). For both datasets, ensemble models were based on two approaches: (i) a quality-weighted mean (MEAN), corresponding to a soft averaging of habitat suitability probabilities weighted by model performance, and (ii) a committee averaging consensus (CA), defined as the proportion of models assigning the highest probability to each class (voting-based classification).

We reduced overprediction by integrating species occurrence patterns, expert range extents from IUCN, and habitat suitability outputs from SDMs using a Bayesian combination approach based on the permanence of ratios (Hoareau et al., 2025). This method refines habitat suitability estimates using prior information on species distributions derived from occurrences and expert range maps, adjusting predictions according to species presence likelihood. For invertebrates, where IUCN expert range maps were not available, species ranges were approximated using ecoregions selected based on occurrence patterns and their spatial distribution within regions, while accounting for uneven sampling effort using a reference group of related species with similar sampling patterns (see Supporting Information S2 for details).

### Uncertainty in spatial conservation prioritization

For the prioritization analysis, we used Zonation 5 software (v2.3), a decision-support framework that produces a hierarchical ranking of planning units based on their contribution to biodiversity representation. Zonation generates two main outputs: (i) spatial priority ranking maps, which assign each planning unit a relative priority value based on its contribution to overall conservation value, and (ii) performance curves, which summarise how feature representation changes as increasing proportions of the landscape are retained according to the priority ranking. All prioritizations were run using the default settings, including the marginal loss removal rule CAZ2, with all species weights set to 1.0 and no additional parameters or constraints applied. CAZ2 provides a good balance between overall average coverage and the coverage of features that are difficult or expensive to represent (Moilanen et al., 2022).

We generated prioritization variants for the two independent SDM datasets (vertebrates and invertebrates), using outputs from each individual algorithm and each ensemble method. For every modelling approach, we contrasted unconstrained scenarios (i.e., using raw SDM outputs) with constrained scenarios in which overprediction was corrected. This resulted in a factorial design structured as dataset × modelling approach × overprediction treatment, yielding 22 prioritization variants in total. These variants formed the basis for quantifying uncertainty in both spatial priority rankings and performance curves (Figure 1).

**Figure 1.**
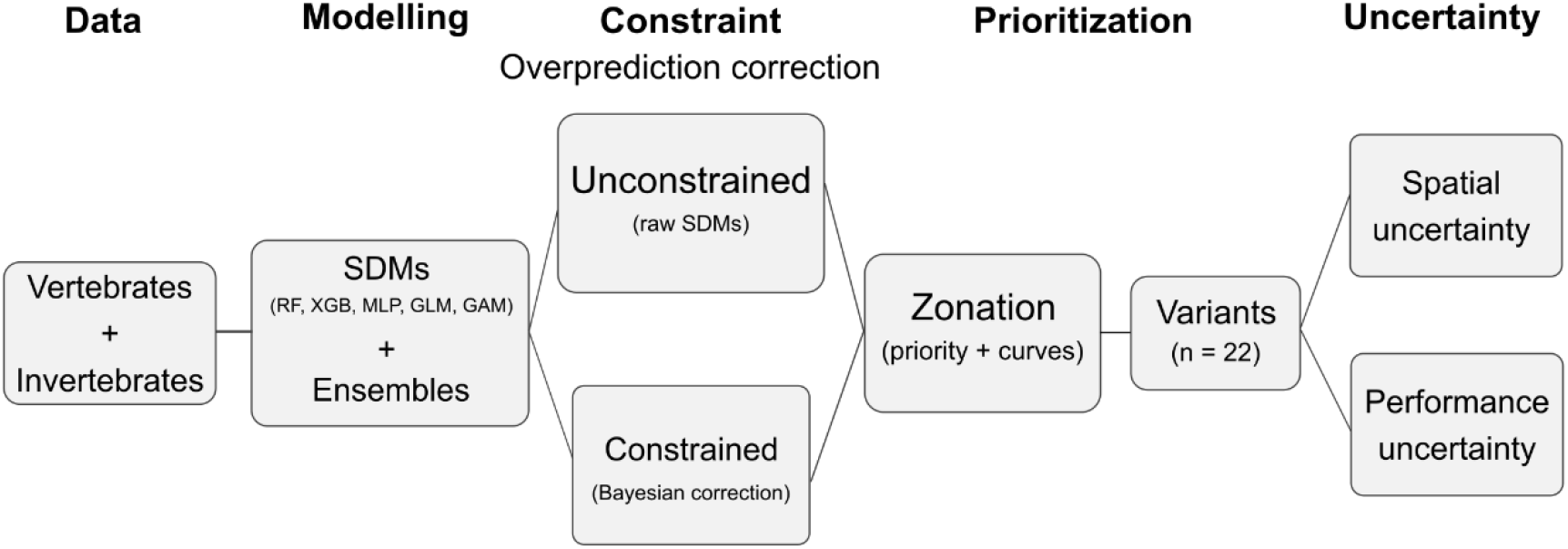
Workflow linking species distribution modelling (SDMs), overprediction correction, and spatial conservation prioritization. SDMs for vertebrates and invertebrates were generated using multiple algorithms and ensemble approaches, and used in both unconstrained and constrained scenarios. Overprediction was reduced using a Bayesian correction framework integrating occurrence data and species range information. These outputs feed into Zonation to produce multiple prioritization variants (n = 22), from which spatial priority rankings and performance curves are derived. Uncertainty is quantified as variation across prioritization outputs, allowing evaluation of how overprediction propagates through modelling choices into conservation prioritization outcomes.

We quantified uncertainty in priority ranking as the pixel-wise standard deviation (SD) across prioritization variants and calculated the difference between unconstrained and constrained scenarios as

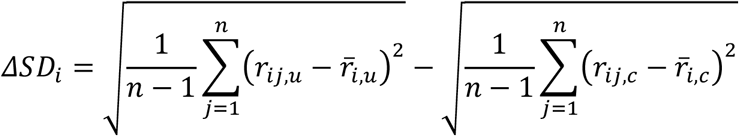

where *r*_*ij,s*_ denotes the prioritization rank assigned by variant *j* to pixel *i* under scenario *s*, 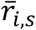 is the mean rank across the *n* variants for pixel *i* under scenario *s*. Here, *u* and *c* refer to the unconstrained and constrained scenarios, respectively. This represents the change in SCP uncertainty for the same planning unit under alternative assumptions. Positive values indicate higher uncertainty under the unconstrained scenario, whereas negative values indicate higher uncertainty under the constrained scenario. We tested differences in prioritization uncertainty between constrained and unconstrained scenarios using repeated subsampling and non-parametric tests (Supporting Information S3).

We also explored which prioritization variants contributed most to scenario-dependent differences in uncertainty. To this end, we applied a paired leave-one-out (PLOO) analysis. In each iteration, the same prioritization variant was removed simultaneously from both constrained and unconstrained sets, and Δ*SD* was recalculated. The contribution of each variant was quantified as the change in ΔSD resulting from its removal compared to the full set of variants (Supporting Information S4).

We applied the same uncertainty framework described above to the performance curves. Uncertainty was quantified as the standard deviation in feature coverage values (i.e., proportion of each species’ distribution retained according to the priority ranking) across prioritization variants, and ΔSD between constrained and unconstrained scenarios was calculated analogously to the spatial formulation. This allowed us to evaluate how uncertainty affects expected conservation performance, allowing direct comparison of uncertainty patterns across the two main prioritization outputs.

## Results

For both vertebrates and invertebrates, the unconstrained scenario showed higher pixel-wise variation across modelling variants compared to the constrained scenario. For vertebrates, median standard deviation (SD) increased by approximately 65% (from 0.048 under the constrained scenario to 0.079 under the unconstrained scenario), while for invertebrates it increased by roughly 55% (from 0.078 under the constrained scenario to 0.122 under the unconstrained scenario) (Figure S1 and S2). The difference in uncertainty between scenarios (ΔSD) was predominantly positive across the landscape. This pattern was consistent across the ten independent random subsamples, with the median ΔSD equal to 0.02 (range 0.02–0.03) for vertebrates and 0.03 (range 0.03–0.04) for invertebrates. These differences were statistically significant, indicating that overprediction consistently amplifies spatial uncertainty in spatial conservation prioritization (Figure 2; Table S1).

**Figure 2.**
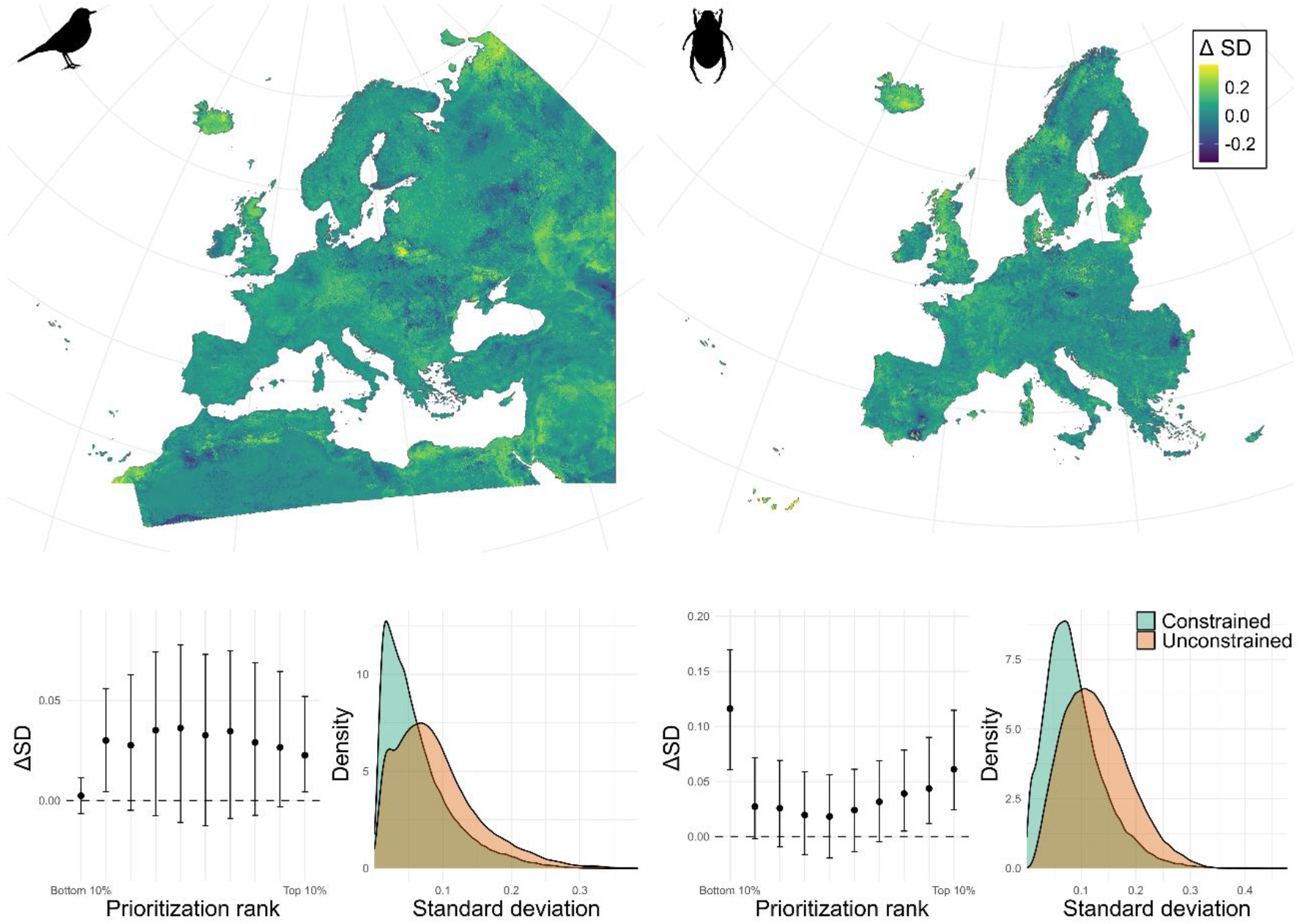
Spatial ranking uncertainty in conservation prioritization outcomes. The top row displays the spatial distribution of ΔSD, where positive values indicate higher uncertainty under the unconstrained scenario and negative values indicate higher uncertainty under the constrained scenario. The bottom row presents two statistical summaries: variation in ΔSD across prioritization rank bins (with interquartile range bars, 25th–75th percentiles) and kernel density estimates of pixel-wise standard deviation (SD) values sampled from constrained and unconstrained scenarios, illustrating the overall distribution of uncertainty.

When summarized by prioritization rank bins, median ΔSD values showed greater variability in intermediate ranks and particularly strong uncertainty reduction in top-priority areas for both datasets, as well as in the lowest ranks (bottom 10%) for the invertebrates. Invertebrates also exhibited lower overall variability across rank bins.

Differences in uncertainty between scenarios (ΔSD) calculated from performance curves reflected the patterns observed in the prioritization ranks. Across all ten subsamples, ΔSD was predominantly positive for vertebrates (median = 0.0618; range = 0.0618–0.0618) and invertebrates (median = 0.0178; range = 0.0178– 0.0178), showing a statistically significant effect. Aggregating by prioritization rank bins, median ΔSD values for vertebrates were consistently positive across all rank intervals. For invertebrates, median ΔSD values were positive across most bins, with the strongest reductions in intermediate ranks, but with slightly negative values in the top-priority bins. For both vertebrates and invertebrates, the constrained scenario consistently led to higher overall performance, as reflected in the performance curves (Figure 3).

**Figure 3.**
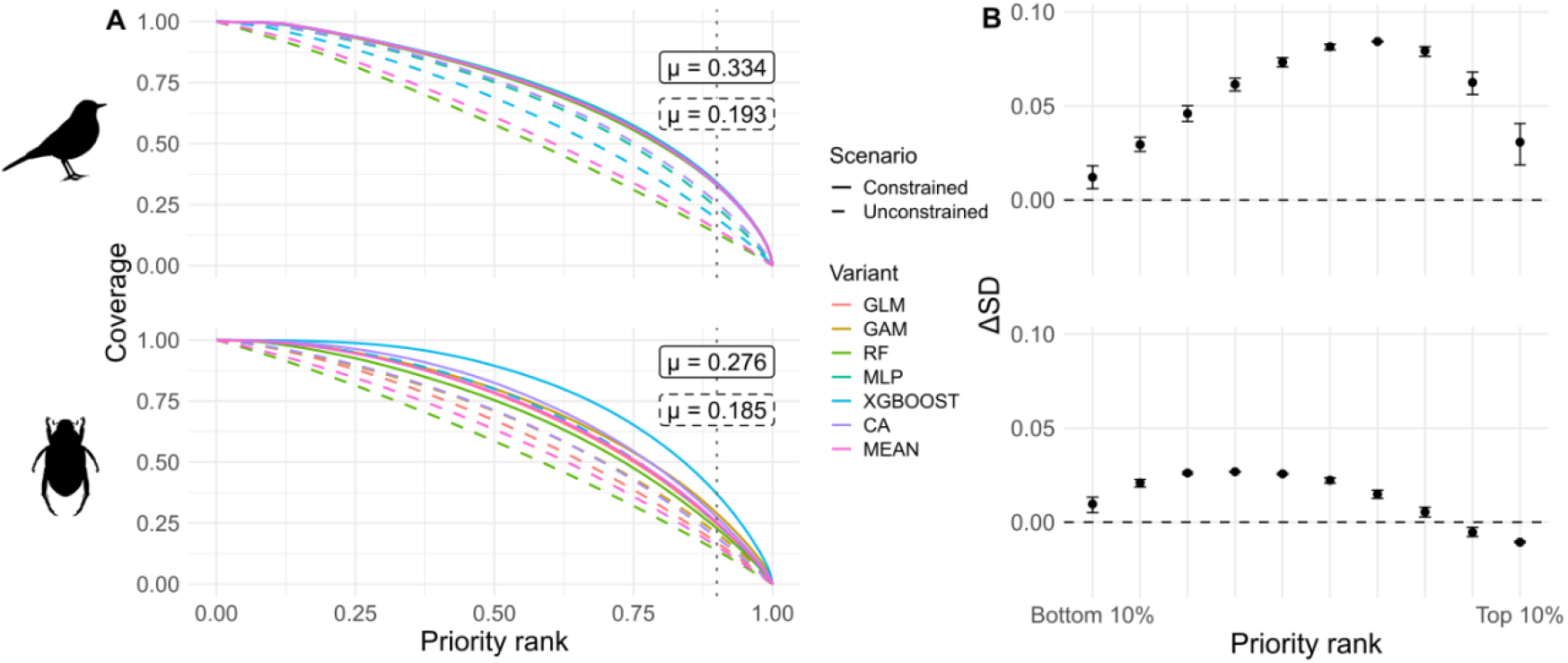
Performance uncertainty in conservation prioritization outcomes. (A) Performance curves showing average coverage for constrained and unconstrained scenarios across prioritization variants. The vertical dotted line highlights the top 10% priority areas, with labels indicating the average coverage (μ) within these areas. Lines are colored by prioritization variant, representing individual SDM algorithms (GLM, GAM, RF, MLP, XGBOOST) and ensemble outputs (CA, MEAN). (B) Variation in ΔSD across prioritization rank intervals. Error bars represent the interquartile range (25th to 75th percentiles).

For the PLOO analysis, we found that some prioritization variants strongly influenced scenario-dependent uncertainty (Figure 4). For spatial ranking uncertainty in vertebrates, RF increased differences between constrained and unconstrained scenarios (+48%), whereas CA reduced the differences between scenarios (−54%). For performance uncertainty in invertebrates, RF also increased differences (+40%).

**Figure 4.**
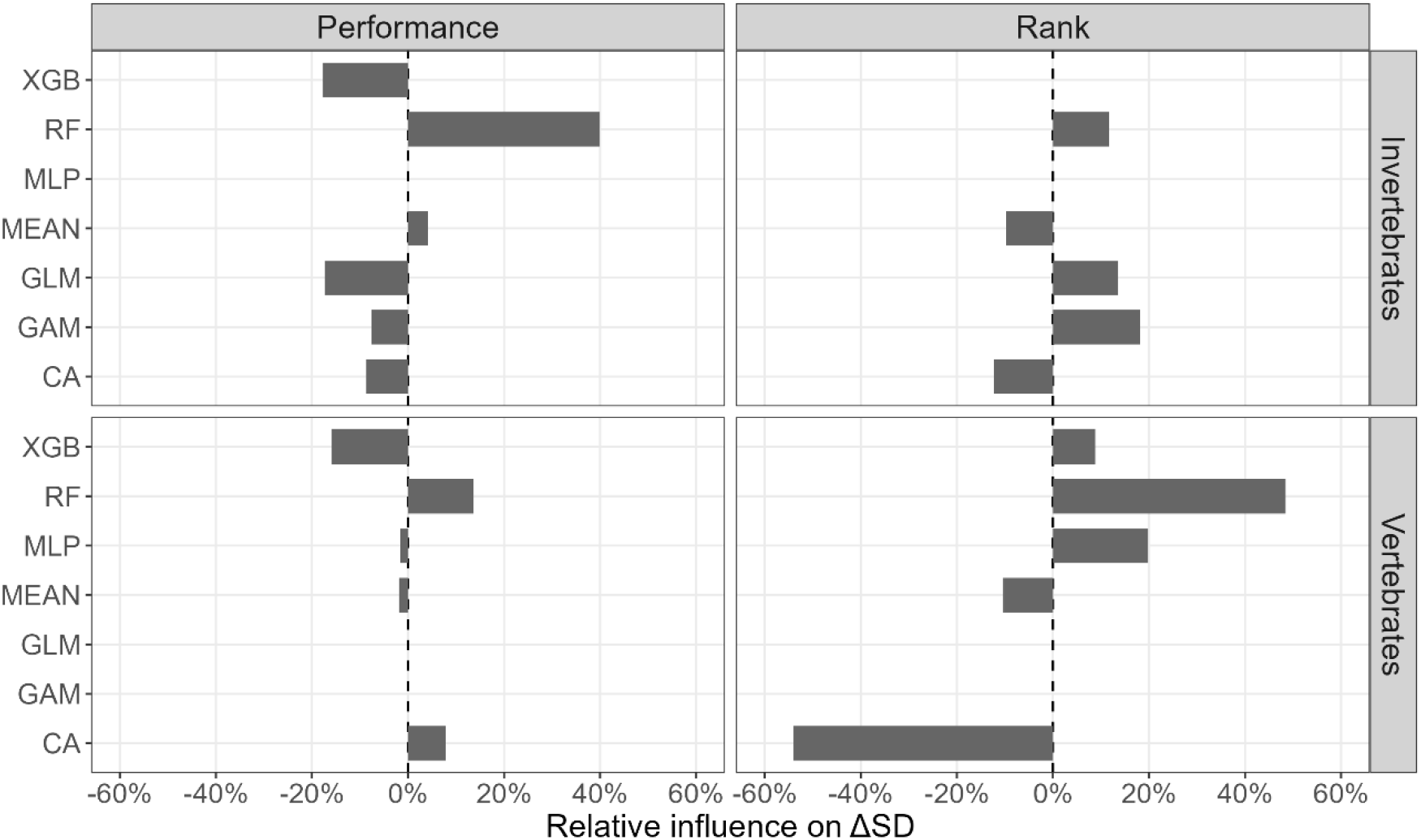
Relative influence of prioritization variants on the median ΔSD, estimated using a paired leave-one-out (PLOO) analysis. Positive values indicate that the variant increases differences in uncertainty between scenarios, whereas negative values indicate that the variant reduces these differences. For example, the RF variant had a relative influence of +48%, meaning that including it results in a larger median ΔSD compared to when it is removed, showing that RF strongly amplifies scenario-dependent uncertainty.

## Discussion

We found that correcting SDM overprediction consistently reduced the propagation of uncertainty in conservation prioritization for both datasets evaluated in this study, with the strongest effects observed in the highest and lowest priority planning units. These areas are particularly important for conservation decision-making, as top-priority units represent sites where protection efforts will have the greatest impact, while bottom-priority units could be suitable for ecological impact avoidance in land-use planning. These findings are consistent with previous studies showing that SDM quality and bias can influence conservation prioritization outcomes (Velazco et al., 2020; Muscatello et al., 2021). We extend these findings by demonstrating the systematic effect of overprediction correction in reducing spatial and performance uncertainty from modelling choices, increasing confidence in prioritization outcomes and supporting more efficient allocation of conservation resources.

Across the landscape, the median spatial priority ranking ΔSD was consistently positive, indicating that overprediction increased uncertainty. Failing to account for overprediction can have significant consequences. Studies have shown that alternative model-building and fitting choices affect outcomes depending on how prioritization is applied, and when SDMs guide the selection of areas for conservation or management actions, these differences can introduce substantial uncertainty, sometimes equivalent to making decisions at random (Muscatello et al., 2021). Overprediction can also distort the spatial allocation of conservation priorities by assigning high conservation value to areas where species may not actually occur (Velazco et al., 2020). In this study, we found that these distortions were accompanied by increased variation in the effect of overprediction in areas where prioritization is more flexible. Intermediate-ranking planning units, for example, exhibited a wider spread of priority ranking ΔSD values, including occasional negative values, reflecting that overprediction correction did not always reduce uncertainty in these areas. This pattern arises because intermediate ranks include many planning units with similar marginal contributions to biodiversity representation, which gives the prioritization algorithm flexibility in how conservation value is allocated across the landscape (Loiselle et al., 2003; Moilanen et al., 2022). In contrast, top-priority units are tightly constrained by their high contribution to biodiversity, and bottom-priority units are constrained by low contribution, leaving little room for variation. These results suggest that while overprediction correction improves overall robustness, intermediate-ranking areas should be interpreted with extra caution, as uncertainty appears more difficult to mitigate in these units.

By analyzing performance curves across prioritization variants, we show that constrained scenarios generally yield improved feature coverage compared to unconstrained scenarios. For vertebrates, improvements were consistent across all rank intervals, while for invertebrates, summarized performance ΔSD values were slightly negative in top-priority bins, indicating that the effect of overprediction correction on performance can vary across rank intervals and datasets. These differences may be related to variation in input accuracy for the constrained approach used in this study. For vertebrates, priors in the Bayesian constraint approach combined SDM outputs with expert range maps and occurrence patterns, providing well-validated distribution boundaries for species and improving the reliability of habitat suitability estimates (Si-Moussi et al., 2025). In contrast, expert range maps were largely unavailable for invertebrates, reflecting the general Wallacean shortfall in invertebrate distributions (Cardoso et al., 2011). Consequently, priors for invertebrates were approximated by selecting ecoregions based on occurrence distributions, potentially leading to greater uncertainty in top-priority areas. Overall, these results emphasize that reducing overprediction improves performance curve reliability, but the extent and pattern of improvements depend on the characteristics and quality of the input data used for overprediction correction.

Based on the PLOO analysis, we found that not all prioritization variants contribute equally to scenario-dependent uncertainty in spatial conservation prioritization. Certain algorithms, such as Random Forest, consistently amplified differences between constrained and unconstrained scenarios, while ensemble approaches tended to reduce these differences. The strong influence of modelling technique choice on spatial predictions and prioritization outcomes has been noted in the literature. Different SDM algorithms produce variable predictions, and ensemble approaches can retain information about this variability, leading to more reliable prioritization results (Meller et al., 2014). Our results highlight that the choice of SDM approach can strongly influence both spatial ranking and performance uncertainties, emphasizing the importance of model-specific contributions when interpreting prioritization outcomes. Moreover, the differential influence of variants reinforces the potential value of ensemble approaches in dampening the impact of overprediction and model variability on prioritization uncertainty. This is particularly important given the sensitivity of conservation planning outcomes to various aspects of species distribution modelling (Lentini and Wintle, 2015; El-Gabbas et al., 2020; Velazco et al., 2020; Muscatello et al., 2021).

Correcting overprediction has clear practical implications for conservation planning. First, we showed that correcting overprediction consistently reduced spatial priority ranking uncertainty, especially in the highest-priority planning units for both vertebrates and invertebrates, highlighting sites where conservation action is likely to have the greatest impact. A similar pattern was observed for the lowest-priority units for invertebrates, indicating that correcting overprediction also increases confidence in areas suitable for impact avoidance. Second, constrained scenarios demonstrated higher overall coverage across prioritization variants, suggesting that conservation policies informed by corrected SDMs are more likely to meet established policy targets for biodiversity representation. Finally, intermediate-ranking planning units exhibited the greatest variability in ΔSD, indicating that overprediction correction does not fully stabilize these areas. Consequently, policies and actions targeting intermediate rank areas should incorporate additional flexibility, monitoring, or adaptive management to account for residual uncertainty in these units.

There are multiple sources of uncertainty in SDMs that can propagate into conservation prioritization outcomes, including algorithm choice, sampling bias, environmental predictors, threshold selection, and model parameterization (Wilson et al., 2005; Dormann et al., 2008; Langford et al., 2009; Merow et al., 2014; El-Gabbas et al., 2020; Velazco et al., 2020; Muscatello et al., 2021). In this study, we specifically focused on overprediction, which we show has a substantial impact on spatial and performance uncertainty. While correcting overprediction consistently reduced uncertainty in our analyses, it is important to emphasize that this is not a universal solution. Other sources of uncertainty warrant further investigation, and different taxa or datasets may respond differently to correction approaches. Despite the insights provided by our study, several limitations should be acknowledged. First, while overprediction correction consistently reduced spatial and performance uncertainty, the magnitude of its effect depends on the quality and completeness of input SDMs and priors used in the constraint approach. For vertebrates, expert-informed priors allowed overprediction to be constrained effectively, whereas for invertebrates, constrained priors were approximated using ecoregions and observation density, which may have introduced additional uncertainty that propagated through the prioritization outputs. Second, the PLOO analysis, while informative about the relative influence of individual prioritization variants, does not quantify additive contributions or explain the mechanistic drivers behind these effects. The metric quantifies the proportional change in overall uncertainty when a given algorithm is removed, but it cannot disentangle interactions among algorithms or partition variance across multiple sources of uncertainty. Third, our factorial design considered a limited set of modelling algorithms and ensemble techniques. Broader algorithmic diversity or alternative ensemble strategies could affect both the magnitude and spatial patterns of uncertainty. Future work could extend this framework to additional taxa, alternative SDM methods, and hierarchical or multi-scale prioritization scenarios, and explore more robust methods to disentangle algorithm-specific versus interaction effects in uncertainty propagation.

Overall, our results highlight that, while no single modelling adjustment can fully resolve the propagation of uncertainty from SDMs to conservation prioritization, overprediction correction consistently has a large effect, systematically reducing uncertainty across datasets, taxa, and modelling approaches. This underscores that applying such corrections is not only beneficial but essential for producing robust and reliable prioritization outcomes.

## Acknowledgments

This research was supported by the NaturaConnect project, funded by the European Union’s Horizon Europe research and innovation programme (grant agreement No. 101060429), and by the Kone Foundation. The authors acknowledge CSC – IT Center for Science, Finland, for providing access to high-performance computing resources.

## References

Beale, C. M. and Lennon, J. J. 2012. Incorporating uncertainty in predictive species distribution modelling. Philosophical Transactions of the Royal Society B: Biological Sciences, 367, 247–258.

Cardoso, P., Erwin, T. L., Borges, P. A. and New, T. R. 2011. The seven impediments in invertebrate conservation and how to overcome them. Biological conservation, 144, 2647–2655.

Diniz-Filho, J. A. F., Loyola, R. D., Raia, P., Mooers, A. O. and Bini, L. M. 2013. Darwinian shortfalls in biodiversity conservation. Trends in Ecology & Evolution, 28, 689–695.

Dormann, C. F., Purschke, O., Marquez, J. R. G., Lautenbach, S. and Schroeder, B. 2008. Components of uncertainty in species distribution analysis: a case study of the great grey shrike. Ecology, 89, 3371–3386.

El-Gabbas, A., Gilbert, F. and Dormann, C. F. 2020. Spatial conservation prioritisation in data-poor countries: a quantitative sensitivity analysis using multiple taxa. Bmc Ecology, 20, 35.

Giakoumi, S. et al. 2025. Advances in systematic conservation planning to meet global biodiversity goals. Trends in Ecology and Evolution, 40, 395–410. 10.1016/j.tree.2024.12.002

Guisan, A., Tingley, R., Baumgartner, J. B., Naujokaitis-Lewis, I., Sutcliffe, P. R., Tulloch, A. I., Regan, T. J., Brotons, L., McDonald-Madden, E. and Mantyka-Pringle, C. 2013. Predicting species distributions for conservation decisions. Ecology letters, 16, 1424–1435.

Hannemann, H., Willis, K. J. and Macias-Fauria, M. 2016. The devil is in the detail: unstable response functions in species distribution models challenge bulk ensemble modelling. Global Ecology and Biogeography, 25, 26–35.

Hoareau, M., Si-Moussi, S. and Thuillier, W. 2025. A Bayesian framework to spatially constrain habitat suitability maps from species distribution models. Authorea Preprint. 10.22541/au.176061482.24539650/v1

Langford, W. T., Gordon, A. and Bastin, L. 2009. When do conservation planning methods deliver? Quantifying the consequences of uncertainty. Ecological Informatics, 4, 123–135.

Lentini, P. E. and Wintle, B. A. 2015. Spatial conservation priorities are highly sensitive to choice of biodiversity surrogates and species distribution model type. Ecography, 38, 1101–1111.

Loiselle, B. A., Howell, C. A., Graham, C. H., Goerck, J. M., Brooks, T., Smith, K. G. and Williams, P. H. 2003. Avoiding Pitfalls of Using Species Distribution Models in Conservation Planning. Conservation Biology, 17, 1591–1600. 10.1111/j.1523-1739.2003.00233.x

Margules, C. R. and Pressey, R. L. 2000. Systematic conservation planning. Nature, 405, 243–253.

Meller, L., Cabeza, M., Pironon, S., Barbet-Massin, M., Maiorano, L., Georges, D. and Thuiller, W. 2014. Ensemble distribution models in conservation prioritization: from consensus predictions to consensus reserve networks. Divers Distrib, 20, 309–321. 10.1111/ddi.12162

Mendes, P., Velazco, S. J. E., de Andrade, A. F. A. and Júnior, P. D. M. 2020. Dealing with overprediction in species distribution models: How adding distance constraints can improve model accuracy. Ecological Modelling, 431, 109180.

Merow, C., Smith, M. J., Edwards Jr, T. C., Guisan, A., McMahon, S. M., Normand, S., Thuiller, W., Wüest, R. O., Zimmermann, N. E. and Elith, J. 2014. What do we gain from simplicity versus complexity in species distribution models? Ecography, 37, 1267–1281.

Moilanen, A. 2012. Spatial conservation prioritization in data-poor areas of the world. Natureza & Conservacao, 10, 1219.

Moilanen, A., Lehtinen, P., Kohonen, I., Jalkanen, J., Virtanen, E. A. and Kujala, H. 2022. Novel methods for spatial prioritization with applications in conservation, land use planning and ecological impact avoidance. Methods in Ecology and Evolution, 13, 1062–1072. 10.1111/2041-210X.13819

Muscatello, A., Elith, J. and Kujala, H. 2021. How decisions about fitting species distribution models affect conservation outcomes. Conservation Biology, 35, 1309–1320.

Si-Moussi, S., Tzivanopoulos, M., Deschamps, G., Hoareau, M., Renaud, J., Lemaire-Patin, R. and Thuiller, W. 2025. Final species and habitat distributions for current and future state. ARPHA Preprints, 6, ARPHA Preprints.

Soberon, J. and Peterson, A. T. 2005. Interpretation of models of fundamental ecological niches and species’ distributional areas.

Thuiller, W., Guéguen, M., Renaud, J., Karger, D. N. and Zimmermann, N. E. 2019. Uncertainty in ensembles of global biodiversity scenarios. Nature communications, 10, 1446.

Thuiller, W. 2024. Ecological niche modelling. Current Biology, 34, R225–R229. 10.1016/j.cub.2024.02.018

Velazco, S. J. E., Ribeiro, B. R., Laureto, L. M. O. and Júnior, P. D. M. 2020. Overprediction of species distribution models in conservation planning: A still neglected issue with strong effects. Biological Conservation, 252, 108822.

Whittaker, R. J., Araújo, M. B., Jepson, P., Ladle, R. J., Watson, J. E. and Willis, K. J. 2005. Conservation biogeography: assessment and prospect. Diversity and distributions, 11, 3–23.

Wilson, K. A., Westphal, M. I., Possingham, H. P. and Elith, J. 2005. Sensitivity of conservation planning to different approaches to using predicted species distribution data. Biological Conservation, 122, 99–112.

Wilson, K. A., Carwardine, J. and Possingham, H. P. 2009. Setting conservation priorities. Annals of the New York Academy of Sciences, 1162, 237–264.

