## Supplementary materials for "Correcting overprediction reduces the propagation of uncertainty from species distribution models into spatial conservation prioritization"

**Affiliations:**
[1] Finnish Museum of Natural History, University of Helsinki, Helsinki, Finland
[2] Univ. Grenoble Alpes, Univ. Savoie Mont Blanc, CNRS, LECA, Grenoble, France

*Supporting Information S1. Species distribution modelling datasets*

Species distribution models (SDMs) for vertebrates and invertebrates were developed at 1 km$^2$ spatial resolution across Europe and ecologically connected surrounding regions (e.g., North Africa, Middle East, and western Russia) to better capture species climatic niches and reduce edge effects. Environmental predictors were selected to represent major climatic, hydrological, edaphic, geological, and land-use gradients influencing species distributions while minimizing collinearity among variables (correlation < 0.80). For both vertebrate and invertebrate datasets, climatic predictors were derived from the CHELSA database (Brun et al., 2022) and included variables related to temperature, precipitation, seasonality, and snow dynamics, such as annual mean temperature, temperature seasonality, annual precipitation, precipitation seasonality, snow cover days, and snow water equivalent. Soil predictors from SoilGrids (Poggio et al., 2021) included soil pH, cation exchange capacity, soil organic carbon, and soil texture variables (sand and silt fractions). Land-use predictors captured information on land cover categories and land-use intensity and were obtained from Sandström et al. (2023) and GlobCover. For vertebrates, which had on average a higher species observation data availability, geological classes and hydrological predictors including coastal proximity, river density, water bodies, wetlands, and water salinity, were additionally used.

For vertebrates, occurrence data were primarily obtained from GBIF and complemented with expert range maps from IUCN and BirdLife. Invertebrate occurrence records were compiled from GBIF and additional expert and citizen-science databases  [e.g., observation.org, SafeGuard (Sentil et al., 2026), and Edaphobase (Burkhardt et al., 2014)]. Vertebrate species distributions were modelled using Random Forest (RF), extreme gradient boosting (XGBoost), and multi-layer perceptron neural networks (MLP). For invertebrates, MLP models were replaced with  generalized linear models (GLM) and generalized additive models (GAM) due to the lower presence data availability. Pseudo-absences were generated using a target group approach where a higher number is selected in areas with higher sampling effort. Additionally, certain absences could be estimated for vertebrates outside of the expert range. For both datasets, SDMs were evaluated using spatial block cross-validation (Valavi et al., 2019) and ensemble predictions retained only models with True Skill Statistic (TSS) ≥ 0.4. Further details on data processing, predictor selection, model calibration, validation, and ensemble construction are provided in Si-Moussi et al. (2025).

*Supporting Information S2. Detailed methods for overprediction correction.*

We reduced overprediction by accounting for various sources of information: occurrence patterns, expert range extents from IUCN and habitat suitability maps from SDM outputs. We used a Bayesian combination method called *permanence of ratios* to improve classical habitat suitability maps with priors (initial knowledge about species distributions) designed from occurrences and expert range maps (see Hoareau et al., 2025 for methodological details). The presence prior is constructed by aggregating occurrences using a spatially-varying kernel density estimation that accounts for local sampling effort, and is transformed to reflect presence likelihood. This map is used together with expert-based maps to construct prior for presence. Presence is determined when there is high occurrence density inside the expert-defined range in an environmentally suitable location, while absence is determined when there are no occurrences in a location outside the expert-defined range and unsuitable for the species. The level of confidence given in intermediate cases is given by the *permanence of ratios* combination formula.

When expert-based range maps were not available, as was the case for the invertebrate dataset in this study, species-specific ranges were constructed using a selection of ecoregions. Ecoregions were relatively selected with respect to each other based on the product of two measures: the observation anomaly and representativity. Observation anomaly was the ratio between the observed number of occurrences inside an ecoregion and the expected number of occurrences this ecoregion would have if the focal species occurrences were distributed among ecoregions similarly to a reference taxon. The reference taxon comprised all occurrences of a broader group of species that included the focal species and that displayed similar sampling patterns. For instance, all occurrences of the Lepidoptera order could be used to correct for spatial sampling bias at the European scale for a focal butterfly species. Depending on the species considered, the reference taxon was chosen at family, order, or a lower taxonomic level, depending on the amount of available occurrence data. Representativity was defined as the percentage of area A of a given ecoregion that was covered by disks of area A/n, centered on the n occurrences within the ecoregion. Representativity was a measure of the spatial spread of occurrences within the ecoregion and therefore reflected the confidence in selecting the entire area without increasing spatial overprediction. Observation anomaly, in turn, measured the strength of the presence signal an ecoregion carried relative to spatial sampling variability.

*Supporting Information S3. Statistical testing of uncertainty differences*

We tested the differences in prioritization uncertainty between constrained and unconstrained scenarios using the distribution of $\Delta SD$ values. Because raster datasets contain a very large number of spatially autocorrelated pixels, statistical testing was performed on repeated random subsamples rather than on the full set of pixels. For each replicate, a random subset of planning units (N = 50000) was drawn without replacement from the study area, and a one-sample Wilcoxon signed-rank test was applied to assess whether the median $\Delta SD$ differed from zero. This procedure was repeated ten times to evaluate the consistency of results across independent subsamples (Table S1).

Table S1. Results of Wilcoxon signed-rank tests evaluating whether the difference in prioritization uncertainty (ΔSD) between scenarios differs from zero across ten random subsamples of planning units.

| Dataset | Replicate | Median ΔSD | p-value |
|---|---:|---:|---|
| Vertebrates | 1 | 0.0249 | <0.001 |
| Vertebrates | 2 | 0.0248 | <0.001 |
| Vertebrates | 3 | 0.0241 | <0.001 |
| Vertebrates | 4 | 0.0252 | <0.001 |
| Vertebrates | 5 | 0.0247 | <0.001 |
| Vertebrates | 6 | 0.0250 | <0.001 |
| Vertebrates | 7 | 0.0245 | <0.001 |
| Vertebrates | 8 | 0.0251 | <0.001 |
| Vertebrates | 9 | 0.0249 | <0.001 |
| Vertebrates | 10 | 0.0250 | <0.001 |
| Invertebrates | 1 | 0.0349 | <0.001 |
| Invertebrates | 2 | 0.0350 | <0.001 |
| Invertebrates | 3 | 0.0349 | <0.001 |
| Invertebrates | 4 | 0.0348 | <0.001 |
| Invertebrates | 5 | 0.0346 | <0.001 |
| Invertebrates | 6 | 0.0348 | <0.001 |
| Invertebrates | 7 | 0.0350 | <0.001 |
| Invertebrates | 8 | 0.0349 | <0.001 |

| Dataset | Replicate | Median ΔSD | p-value |
|---|---|---|---|
| Invertebrates | 9 | 0.0349 | <0.001 |
| Invertebrates | 10 | 0.0351 | <0.001 |

*Supporting Information S4. Paired leave-one-out (PLOO) influence metric*

We used a paired leave-one-out (PLOO) approach to quantify the relative contribution of each modelling variant to scenario-dependent uncertainty. Each variant was sequentially excluded from both constrained and unconstrained prioritization sets. To facilitate comparison across variants, we expressed this influence as a relative percentage of the full-model median ΔSD, with $j$ denoting the prioritization variant removed in each iteration:

$$Relative\ influence_j = \frac{\mathrm{Median}(\Delta SD_{full}) - \mathrm{Median}(\Delta SD_{-j})}{\mathrm{Median}(\Delta SD_{full})} \times 100$$

This metric quantifies the proportional change in the overall median ΔSD when a given algorithm is removed. Positive values indicate that the variant increases differences in SCP uncertainty between scenarios, whereas negative values indicate that the variant reduces these differences. Note that this metric reflects the sensitivity of the median ΔSD to removal of each variant and does not represent variance partitioning or explained contribution in an additive sense.

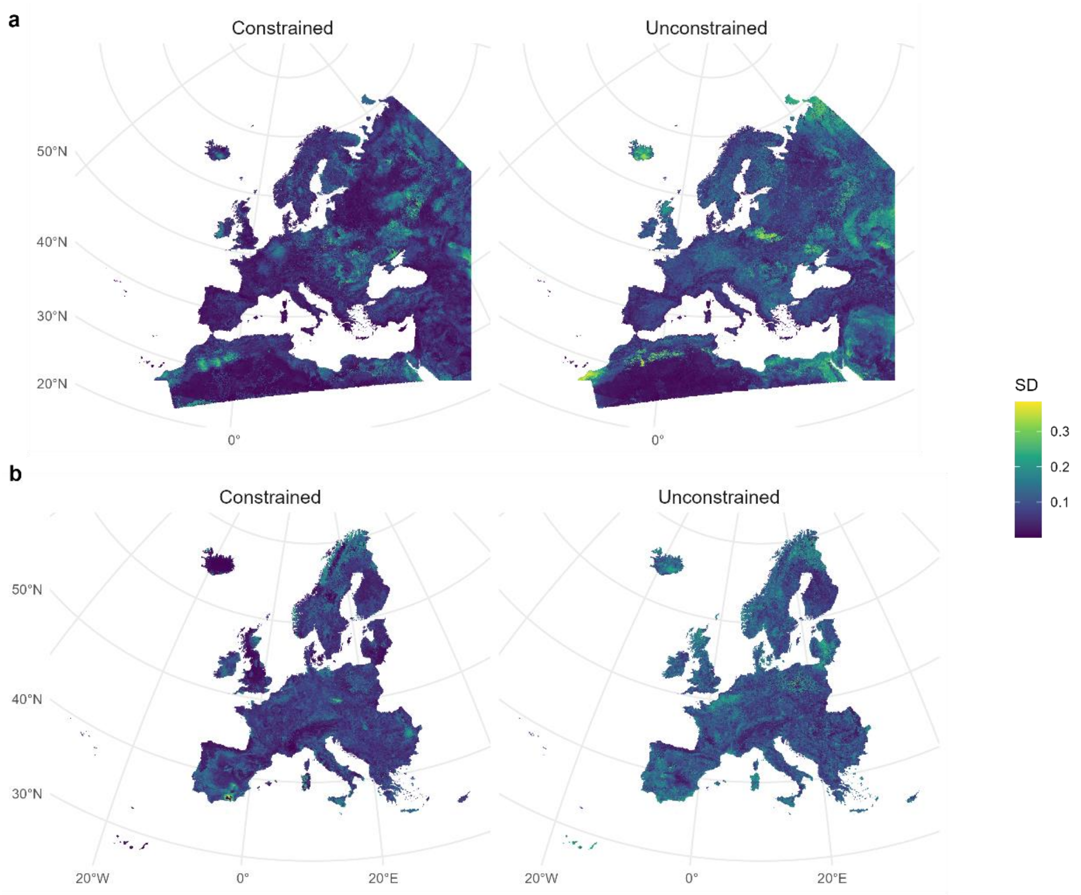


Figure S1. Pixel-wise standard deviation (SD) across prioritization variants under constrained and unconstrained scenarios. Each row shows one taxonomic group: vertebrates (a) and invertebrates (b), with columns representing constrained and unconstrained prioritization.

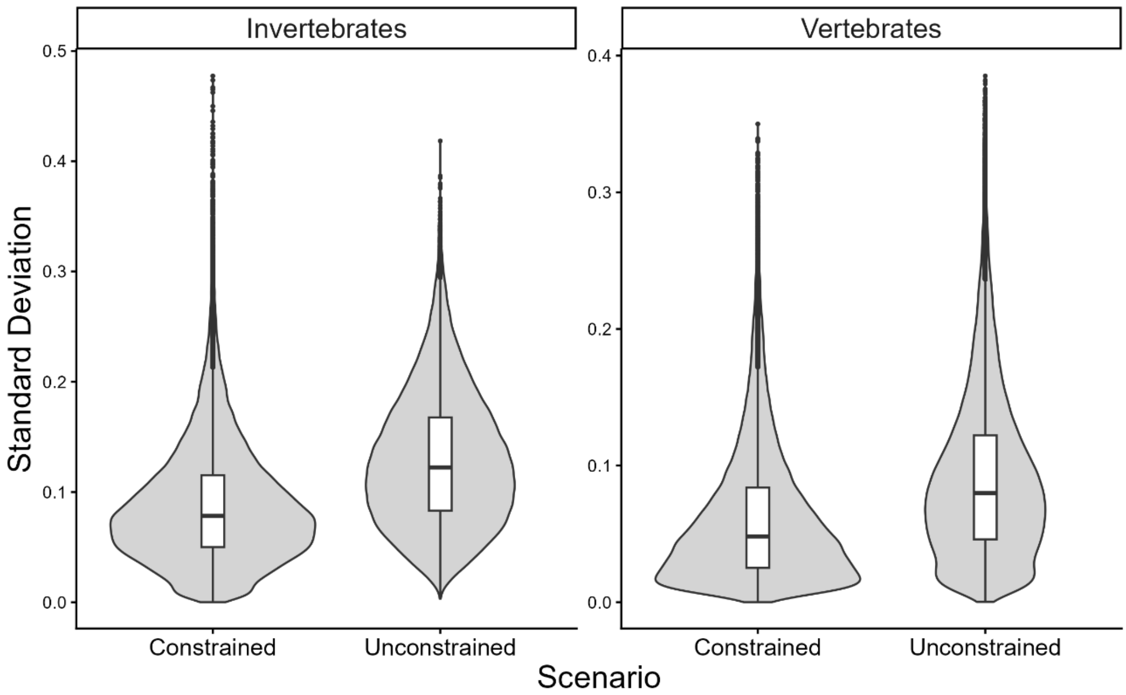

Figure S2. Standard deviation (SD) across prioritization variants for constrained and unconstrained scenarios. Values are based on a random sample of 100,000 planning units per scenario across the study area.

**References**

Brun, P., Zimmermann, N. E., Hari, C., Pellissier, L. and Karger, D. N. 2022. Global climate-related predictors at kilometer resolution for the past and future. *Earth System Science Data,* 14, 5573-5603.

Burkhardt, U., Russell, D., Decker, P., Döhler, M., Höfer, H., Lesch, S., Rick, S., Römbke, J., Trog, C. and Vorwald, J. 2014. The Edaphobase project of GBIF-Germany—A new online soil-zoological data warehouse. *Applied Soil Ecology,* 83, 3-12.

Hoareau, M., Si-Moussi, S. and Thuillier, W. 2025. A Bayesian framework to spatially constrain habitat suitability maps from species distribution models. *Authorea Preprint*. 10.22541/au.176061482.24539650/v1

Poggio, L., De Sousa, L. M., Batjes, N. H., Heuvelink, G., Kempen, B., Ribeiro, E. and Rossiter, D. 2021. SoilGrids 2.0: producing soil information for the globe with quantified spatial uncertainty. *Soil,* 7, 217-240.

Sandström, E., Namasivayam, A., Oostdijk, S., Scherpenhuijzen, N., Debonne, N. and Verburg, P. (2023). Land system map for Europe (V6 ed.): DataverseNL.

Sentil, A., Miličić, M., Benrezkallah, J., Ačanski, J., Andrić, A., Aubert, M., Bartomeus, I., Biella, P., Boustani, M. and Carstensen, L. B. 2026. Synthesised database of wild bee and hoverfly records in Europe. *Scientific Data*.

Si-Moussi, S., Tzivanopoulos, M., Deschamps, G., Hoareau, M., Renaud, J., Lemaire-Patin, R. and Thuiller, W. 2025. Final species and habitat distributions for current and future state. *ARPHA Preprints,* 6, ARPHA Preprints.

Valavi, R., Elith, J., Lahoz-Monfort, J. J. and Guillera-Arroita, G. 2019. blockCV: An r package for generating spatially or environmentally separated folds for k-fold cross-validation of species distribution models. *Methods in Ecology and Evolution,* 10, 225-232. https://doi.org/10.1111/2041-210X.13107
